# DeltaNeTS+: Elucidating the mechanism of drugs and diseases using gene expression and transcriptional regulatory networks

**DOI:** 10.1101/788968

**Authors:** Heeju Noh, Ziyi Hua, Panagiotis Chrysinas, Jason E. Shoemaker, Rudiyanto Gunawan

## Abstract

**Background:** Knowledge on the molecular targets of diseases and drugs is crucial for elucidating disease pathogenesis and mechanism of action of drugs, and for driving drug discovery and treatment formulation. In this regard, high-throughput gene transcriptional profiling has become a leading technology, generating whole-genome data on the transcriptional alterations caused by diseases or drug compounds. However, identifying direct gene targets, especially in the background of indirect (downstream) effects, based on differential gene expressions is difficult due to the complexity of gene regulatory network governing the gene transcriptional processes.

**Results:** In this work, we developed a network analysis method, called DeltaNeTS+, for inferring direct gene targets of drugs and diseases from gene transcriptional profiles. DeltaNeTS+ relies on a gene regulatory network model to identify direct perturbations to the transcription of genes. Importantly, DeltaNeTS+ is able to combine both steady-state and time-course gene expression profiles, as well as to leverage information on the gene network structure that is increasingly becoming available for a multitude of organisms, including human. We demonstrated the power of DeltaNeTS+ in predicting gene targets using gene expression data in complex organisms, including *Caenorhabditis elegans* and human cell lines (T-cell and Calu-3). More specifically, in an application to time-course gene expression profiles of influenza A H1N1 (swine flu) and H5N1 (avian flu) infection, DeltaNeTS+ shed light on the key differences of dynamic cellular perturbations caused by the two influenza strains.

**Conclusion:** DeltaNeTS+ is an enabling tool to infer gene transcriptional perturbations caused by diseases and drugs from gene transcriptional profiles. By incorporating available information on gene network structure, DeltaNeTS+ produces accurate predictions of direct gene targets from a small sample size (~10s). DeltaNeTS+ can freely downloaded from http://www.github.com/cabsel/deltanetsplus.

## Background

Analyzing the molecular mechanism of drugs and diseases is a central task in drug discovery. For drugs, this task involves identifying the molecular targets whose interaction with the compound is associated with its pharmacological activity. The knowledge of the molecular mechanism of action (MOA) of a drug is important in ascertaining not only the therapeutic efficacy of this drug, but also any potential toxicity and side effects. On the other hand, insights into the molecular mechanism of a disease may lead to a better understanding of its pathogenesis and possible to new and better treatment formulations. In this regard, gene transcriptional profiling has emerged as a viable high-throughput platform for drug discovery and drug target identification [1, 2] and for studying disease mechanism [3]. However, determining the direct molecular targets through which a drug or a disease exert its effects using gene transcriptional profiles remains a major bioinformatics challenge. Changes in the gene expression caused by a drug or a disease may arise directly from the action of the drug or disease, or indirectly as downstream or secondary effects. Delineating direct and indirect gene targets from gene transcriptional profiles is further complicated by the fact that gene expression is a highly regulated process that involves a complex and context-specific gene regulatory network (GRN).

Many strategies have been proposed for inferring gene targets from gene transcriptional data [4]. In general, these strategies fall in two main categories: comparative analysis and network analysis methods. The former involves comparing the transcriptional profiles of interest with a library of reference profiles with known targets [1, 2, 5, 6], under the assumption that a likeness in the transcriptional profiles is indicative of similarity in the molecular targets. The latter class of methods uses a model of GRN to account for gene transcriptional regulations when analyzing gene expression data. A number of network analysis methods, notably causal analysis [7, 8] and DeMAND [9], employs graph models of the GRNs with nodes representing the genes and edges representing gene-gene interactions. Here, gene target identification is typically formulated as a statistical hypothesis test using the gene transcriptional profiles. Another group of network analysis methods, including network identification by multiple regression (NIR) [10], mode of action by network identification (MNI) [11], sparse simultaneous equation model (SSEM) [12], and DeltaNet [13], relies on a mechanistic model of the gene transcriptional regulatory network. By invoking a pseudo-steady state assumption, the inference of gene targets from gene expression data is recasted as a regression problem. Generally speaking, a gene is scored highly as a potential target when its expression deviates significantly from what the GRN model predicts based on the expression of its transcription factors (TFs).

Here, we present a much-improved network analysis method of our previous algorithm DeltaNet [13] to address two shortcomings: analysis of time-series data and incorporation of prior information on the GRN structure. As we have demonstrated earlier [14], DeltaNet performed poorly when using time-series transcriptional data due to the invalidity of the underlying steady state assumption. Such an issue likely applies to similar algorithms employing pseudo steady state assumption, such as NIR, MNI and SSEM. There is a clear need to accommodate time-series expression data, as they constitute an important class of gene expression datasets. A notable example is the Connectivity Map (CMAP), which includes over 1.5 million time-series gene expression profiles from 9 different cell types across roughly 5000 small-molecule compounds and 3000 genetic reagents.

The second shortcoming of any network analysis methods is the uncertainty in the gene network model employed in the analysis. In DeltaNet, as well as in NIR, MNI and SSEM, the GRN is reconstructed from the gene expression data as an intermediate step or implicitly in the inference. But, as we and many others have shown [15, 16], such GRN reconstructions constitute an undetermined problem and thus the reconstructed GRN often has high uncertainty. In parallel, we have seen a tremendous progress over the past decade in the mapping of transcriptional regulatory elements [17], the measurements of promoter/enhancer activity [18], and the identification of TF binding motifs and binding sites [19]. Such information has enabled the reconstruction of GRN graphs for ~300 cell types in human [20]. Thus, there is an obvious opportunity and necessity to combine diverse datasets on the GRN structure with the gene transcriptional data within the network analysis paradigm for gene target inference.

In this work, we developed DeltaNeTS+. DeltaNeTS+ is capable of combining steady-state and time-series gene transcriptional data for gene target scoring. DeltaNeTS+ is further able to incorporate prior information of the structure of the GRN for the target inference. We demonstrated the superiority of DeltaNeTS+ over well-established procedures, including differential expression analysis and TSNI (Time Series Network Identification) [21], using gene transcriptional datasets of complex organisms, including *Caenorhabditis elegans* and human. Notably, the application of DeltaNeTS+ to gene expression datasets from human lung Calu-3 cells infected by influenza strains H5N1 (avian flu) and H1N1 (swine flu) reveals key differences in the cellular responses to the two strains.

## Methods

### DeltaNeTS+

DeltaNeTS+ is a network analysis method for inferring causal gene targets from time series gene expression data. DeltaNeTS+ relies on an ordinary differential equation (ODE) model of gene transcriptional process, as follow: [22]

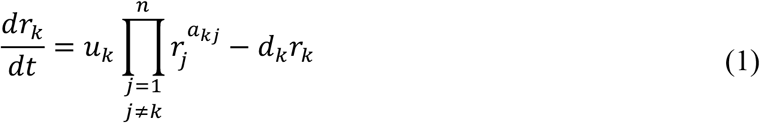

where *r_k_* denotes the concentration of gene *k* transcripts (mRNA), *u_k_* and *d_k_* denote the transcription and degradation rate constants of gene *k*, respectively, *a_kj_* denotes the regulatory control of gene *j* on gene *k*, and *n* is the number of genes. The magnitude and sign of *a_kj_* indicate the strength and mode of the regulatory interaction, respectively. A positive *a_kj_* indicates activation, while a negative *a_kj_* indicates repression. As commonly done, *a_jj_*’s are assumed to be zero (i.e., no self-regulatory loops). Taking pseudo steady state assumption and logarithm transformation, Equation (1) can be simplified into,

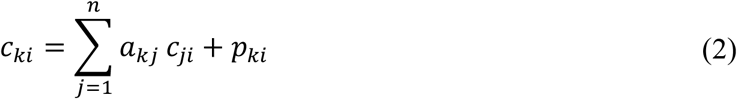

where *c_ki_* = log_2_(*r_ki_* / *r_kb_*) denotes the log-2 fold change (log2FC) of mRNA levels of gene *k* between treatment sample *i* and corresponding control *b*, and *p_ki_* = log((*u_ki_* / *d_ki_*) / (*u_kb_* / *d_kb_*)) denotes the effects of treatment on gene *k* in sample *i*. The variable *p_ki_* captures the perturbations on gene *k* and is the unknown variable of interest. A positive (negative) *p_ki_* indicates that the treatment causes a higher (lower) level of gene *k* transcription than what is expected from the changes in the transcription factors or regulators of the gene.

As we often have more than one samples for a given treatment (e.g., technical repeats, multiple time points), we may assign the same perturbations to all of the related samples. For *M* different treatments among *m* samples (*M*<*m*), the following equation in a matrix-vector format can be considered:

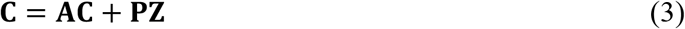

where **C** is the *n×m* matrix of log2FCs of *n* genes in *m* samples, **A** is the *n×n* matrix of *a_kj_*’s describing the GRN, **P** is the *n×M* matrix of perturbation coefficients, and **Z** is an *M×m* {0, 1} mapping matrix that assigns the specific perturbation variable *p* to the appropriate sample. Note that **Z** becomes an *m×m* identity matrix when the perturbations are treated as distinct among the entire *m* samples.

For time-series data, we assume that mRNA concentrations vary between sampling time points such that

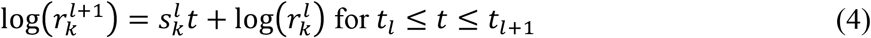

where *t*_*l*_ denotes the *l-*th time point, 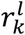 is the mRNA concentration of gene *k* at the *l*-th time point, and 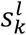 is the slope between the two time points. Using Equation (1), the first order derivative of the logarithm of *r_k_* can be rewritten as the following:

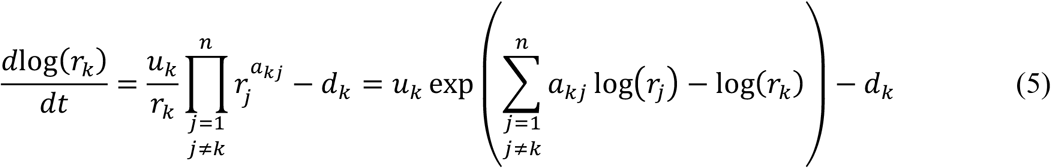

By substituting Equation (4) for *r_k_* in Equation (5), the following equation can be derived for *t*_*l*_ ≤ *t* ≤ *t*_*l*+1_:

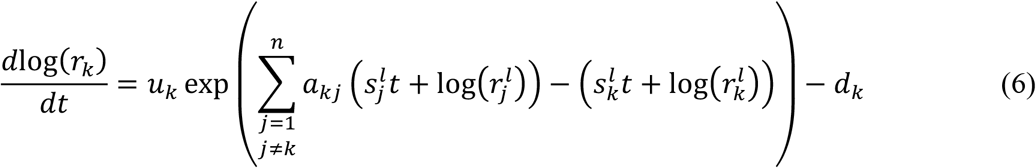

The derivative of Equation (6) (i.e. the second derivative of log(*r_k_*)) is zero under the pseudo steady-state assumption made in Equation (4).

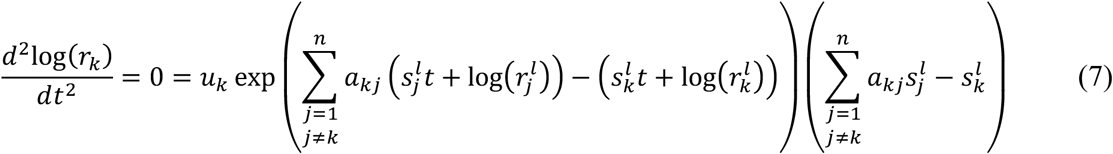

Here, *u_k_* and *d_k_* are assumed to be the same between the *l*-th and *l*+1-th time points, i.e. the perturbations are constant in this time window. Since *u_k_* and the exponential term in Equation (7) are always positive, the remaining term should be zero. Therefore, the following relationship between the variable 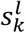 and the network coefficient *a_kj_* can be obtained:

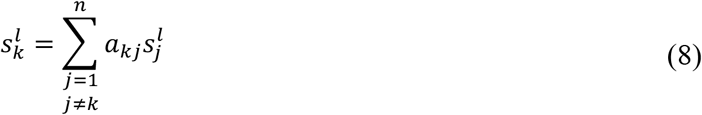

The matrix form of Equation (8) for *n* genes and *m* samples is the same as following:

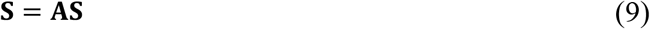

where **S** is the *n×m* matrix of the slopes calculated from time series log2FCs data. In DeltaNeTS+, the slopes of the time series gene expression profiles were calculated using 2^nd^-order accurate finite difference approximations at each sampling time point [23]. For the first and last time points, forward and backward finite difference were used, respectively, while for middle time points, a centered difference approximation was used.

Combining Equation (3) and Equation (9), DeltaNeTS+ calculates the unknown variables **A** and **P** row by row using the following equation:

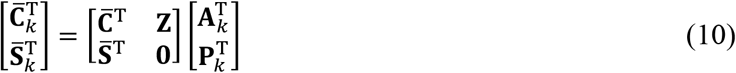

where 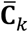, 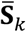, **A**_*k*_, and **P**_*k*_ are the row vectors of the matrices 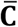, 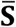, **A**, and **P** for gene *k* and **0** is the *m×M* zero matrix. The matrices 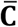 and 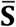 are the normalized log2FC and slope matrices **C** and **S** such that each of the matrices has a 2-norm equals to the square root of the number of samples in the matrix (i.e. the number of columns in the matrix). The normalization is set to balance the contributions from the two matrices in determining **A** and **P**.

In DeltaNeTS+, when information on the structure of the GRN is available – for each gene *k*, we have information on its transcription factors – we can reduce the dimension of the problem stated in Equation (10) by restricting the inference of **A** only for the subset of genes related to the TFs of gene *k*:

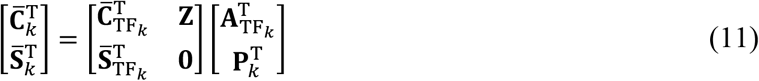

where the subscript TF_*k*_ refers to the subset of genes corresponding to the TFs of gene *k*. Equation (12) is solved using ridge regression [24]. When GRN structure is not provided or not available, DeltaNeTS+ solves Equation (10) by using LASSO regularization [25] to predict a sparse matrix **A**_*k*_ and **P**_*k*_. We implemented LASSO and ridge regression in DeltaNeTS+ using the GLMNET algorithm [26].

### Gene expression data

We applied DeltaNeTS+ to time-series gene expression data of *C. elegans* embryo [27], human cord blood CD4+ T cells [28], and human lung cancer Calu-3 cells [29-31]. For *C. elegans* embryo, log2 intensity data which were normalized by robust multi-array average (RMA) method were obtained from ArrayExpress [32] (accession number: E-GEOD-2180). Log2FC of gene expressions and its statistical significance (Benjamini-Hochberg adjusted *p*-value) between gene knockout and wild-type conditions at each time point was calculated using a linear fit model and empirical Bayes method in the *limma* package of Bioconductor. The probe sets were mapped to the official gene symbols in *celegans.db*. In the case of multiple probe sets mapping to a gene symbol, we take the log2FC from the probe set with the smallest average adjusted *p*-value over the samples.

For human T cells, log2 intensity data by quantile normalization were obtained from ArrayExpress (accession number: E-GEOD-17851). In the same way as *C. elegans*, log2FC values were calculated between gene knockout and wild-type conditions at each time point using *limma*. The probe sets were mapped to the gene symbols from the Illumina human-6 v2.0 expression beadchip data in Gene Expression Omnibus (GEO) [33] (accession number: GPL6102), and for the multiple probe sets mapping to the same gene, the probe set with the smallest average adjusted *p*-value across all samples was chosen.

For Calu-3 data, we compiled the raw *Agilent Whole Human Genome 4×44K* microarray data from GSE33264 for IFN-α and IFN-γ experiments [31] and from GSE37571 and GSE33142 for H1N1 and H5N1 experiments [29, 30]. The raw data were background-corrected and normalized using *normexp* and *quantile* methods in *limma* package of Bioconductor. The log2FCs between virus (or interferon) and mock samples at each time point were calculated using *limma*, with their statistical *p*-value adjusted by Benjamini-Hockberg method. The probe sets were mapped to the official gene symbols in *hgug4112a.db* package. For a gene with multiple probe sets, we chose the data from the probe set with the smallest average adjusted *p*-value.

During preprocessing of time-series data, we substituted any time-series log2FC gene expression data that were not statistically significant with linearly interpolated values using adjacent time points with statistically significant log2FCs. Unless mentioned differently, we used Benjamini-Hochberg adjusted *p*-value < 0.05 to establish statistical significance. If the log2FC values of a gene were not statistically significant at any time point, we set the log2FCs to zero.

### Gene regulatory networks

The GRN for *C. elegans* embryo data set was obtained from TF-gene interactions for *C. elegans* in TF2DNA database [34], where TF binding motifs and their regulated genes were identified by calculating the binding affinity based on the known 3D structure of TF-DNA complexes. The GRN for *C. elegans* was composed of 355,080 edges from 48 TFs to 15,738 genes.

Meanwhile, the GRNs for human T-cell and Calu-3 data sets were obtained from TF-gene interactions specific for human cord blood-derived cells and human epithelium lung cancer cells, respectively, available in Regulatory Circuit database [20]. We only used TF-gene interactions with a confidence score greater than 0.1. The GRNs for human T-cells and Calu-3 consisted of 11,955 edges pointing from 438 TFs to 2,385 genes and 42,145 edges pointing from 515 TFs to 7,125 genes, respectively.

### Performance assessment in knocked-down target predictions

For performance evaluation of DeltaNeTS+ in predicting knocked-down gene targets, we calculated the area under receiver operating characteristic curve (AUROC) and the area under precision and recall curve (AUPR) for each sample, using PRROC package in R. For the AUROC and AUPR calculation, the absolute values in *p_ki_* were used as a threshold.

### Enrichment analysis of calu-3 data

For interferon case study, the top 50 genes ranked by each method (DeltaNeTS+, TSNI, and log2FC) were used for Gene Ontology (GO) and Reactome pathway enrichment analysis using Enrichr [35]. For influenza A viral infection case study, the averaged **P** values of DeltaNeTS+ for each phase (phase 1: 0 to 7 hours, phase 2: 7 to 18 hours, phase 3: >18 hours) were used for the gene set enrichment analysis (GSEA) of Reactome pathways[36] using ReactomePA package[37] in R. Before the enrichment analysis, genes were sorted based on the average **P** for each time phase, and genes with no perturbation score were excluded during the GSEA. Afterwards, the significance score (−log_10_ p-value) of the enriched pathways was calculated among the pathways with positive enrichment score from GSEA. The illustration of the results only shows the highest level of pathway information in the Reactome hierarchy (https://reactome.org/PathwayBrowser), while the significance score is the best significance score (highest −log_10_ *p*-value) of the sub-pathways.

### Weighted Gene Co-expression Network Analysis (WGCNA)

WGCNA [38] was applied to the DeltaNeTS+ score of both H1N1 (influenza A/CA/04/09) and H5N1 (influenza A/VN/1203/04) samples. The H1N1 data were from 9 time points (0, 3, 7, 12, 18, 24, 30, 36, and 48 hours) and the H5N1 data were from 6 time points (0, 3, 7, 12, 18, and 24 hours). In WGCNA analysis, we computed modules with a minimum of 200 genes using the signed network option (soft-thresholding power=18). Afterwards, GO and Reactome enrichment analysis were also performed for each module using ReactomePA and clusterProfiler packages in R.

## Results

### Predicting genetic perturbations

We tested the performance of DeltaNeTS+ by inferring gene perturbations from time series *Caenorhabditis elegans* and human T-cell gene expression profiles from RNA interference (RNAi) experiments. The *C. elegans* dataset comprises three genetic perturbation experiments [27], each of which provides gene expression data across 10 time points after *skn*-1 and *pal*-1 knockdowns by shRNA on *mex*-3 or *pie*-1 mutated cells. The human T-cell dataset includes time-series gene expression data measured over 4 time points after STAT6 knockdown by shRNA [28] (see Table 1). When applying DeltaNeTS+ on these data, gene regulatory network graphs for *C. elegans* and human T-cells were used as prior information (see Methods).

**Table 1.**
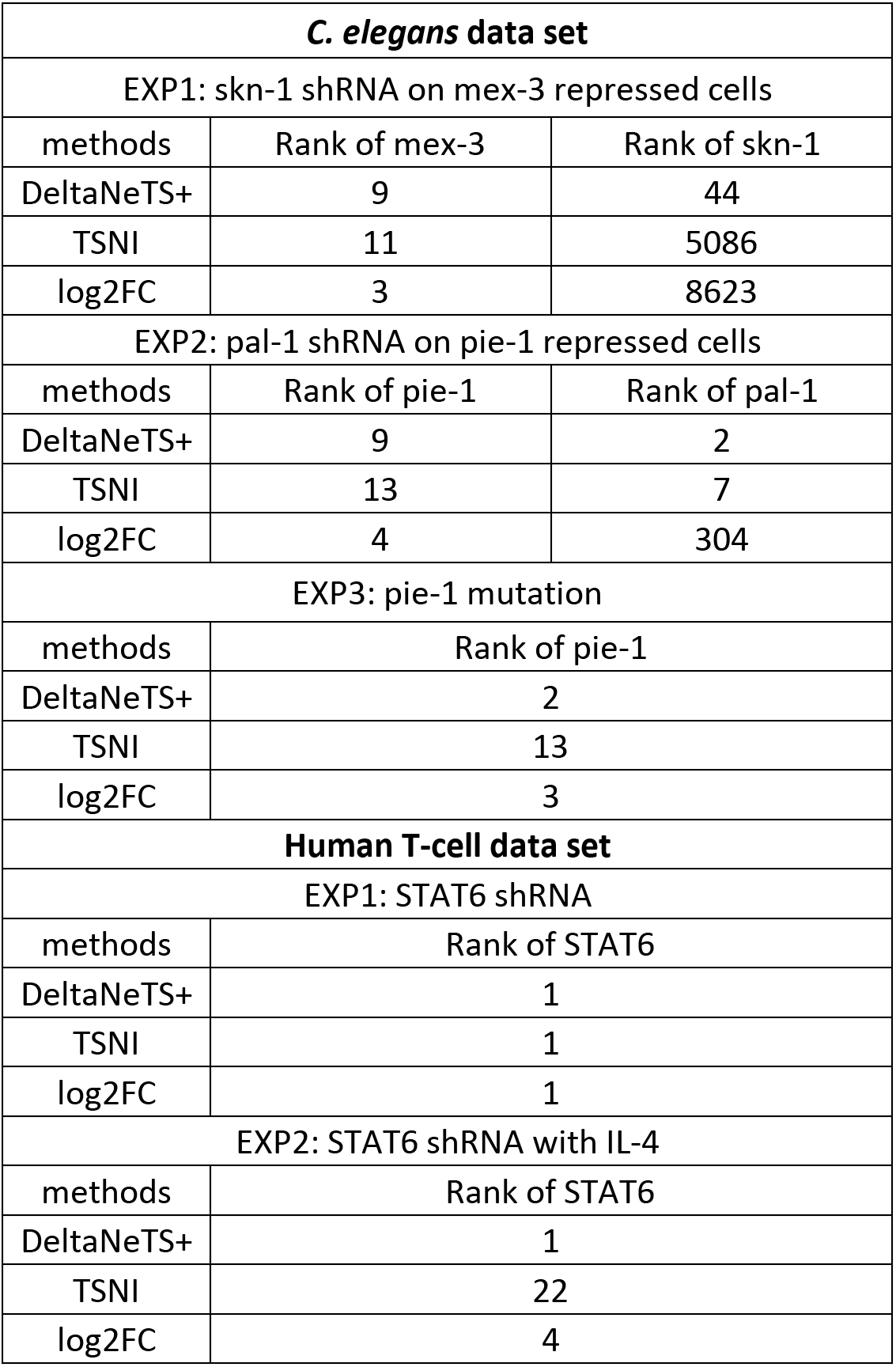
Rank prediction of gene targets in *C. elegans* and human T-cell data sets by DeltaNeTS+, TSNI, and log2FC magnitudes. In TSNI implementation, the first principle component (#PC=1) was used for the analysis of both data sets.

| *C. elegans* data set |  |  |
| --- | --- | --- |
| EXP1: skn-1 shRNA on mex-3 repressed cells |  |  |
| methods | Rank of mex-3 | Rank of skn-1 |
| DeltaNeTS+ | 9 | 44 |
| TSNI | 11 | 5086 |
| log2FC | 3 | 8623 |
| EXP2: pal-1 shRNA on pie-1 repressed cells |  |  |
| methods | Rank of pie-1 | Rank of pal-1 |
| DeltaNeTS+ | 9 | 2 |
| TSNI | 13 | 7 |
| log2FC | 4 | 304 |
| EXP3: pie-1 mutation |  |  |
| methods | Rank of pie-1 |  |
| DeltaNeTS+ | 2 |  |
| TSNI | 13 |  |
| log2FC | 3 |  |
| Human T-cell data set |  |  |
| EXP1: STAT6 shRNA |  |  |
| methods | Rank of STAT6 |  |
| DeltaNeTS+ | 1 |  |
| TSNI | 1 |  |
| log2FC | 1 |  |
| EXP2: STAT6 shRNA with IL-4 |  |  |
| methods | Rank of STAT6 |  |
| DeltaNeTS+ | 1 |  |
| TSNI | 22 |  |
| log2FC | 4 |  |

For comparison purposes, we also generated gene perturbation predictions based on the magnitudes of log2FC values and using TSNI [21]. For log2FC analysis, the gene target candidates were ranked in decreasing order of the absolute value of the log2FCs. For TSNI, we used the first principal component to generate the gene target predictions as this setting gave the best accuracy among the trials using principle components between 1 and 3. We also compared the accuracy of the gene target predictions under two different scenarios; the first is where the gene perturbations are time invariant, and the other is where the gene perturbations are allowed to vary over time. Note that we were unable to perform TSNI for time-varying perturbations, as the algorithm is unable to consider such perturbations.

Table 1 shows the ranking of the genes targeted by shRNA or mutation in *C. elegans* and human T-cell data with the gene perturbations set to be time-invariant, i.e. the gene perturbations are the same for all samples from the same treatment/condition. Except for *skn*-1, DeltaNeTS+ placed the known targets (*mex*-3, *pal*-1, *pie*-1, and STAT6) among the top 10 of the candidate gene targets. Notably, in almost all instances, DeltaNeTS+ ranked the known targets higher than log2FC analysis and TSNI. The transcription factor *skn*-1 regulates critical biological pathways related to oxidative stress responses and lifespan of *C. elegans* [39]. Silencing *skn*-1 alters the transcription of a large number of genes (>10,000), which complicates the identification of the true gene perturbation in these samples. Nevertheless, DeltaNeTS+ was able to put *skn*-1 at a much higher rank than log2FC and TSNI.

When using time-varying perturbation (i.e. the targets can vary across different time samples of the same treatment/condition), DeltaNeTS+ again performed better than only looking at differential expression (log2FC), as illustrated in Figure 1. The log2FC of samples from the early time points actually gave a reasonably accurate indication of the direct gene perturbations (see Supplementary Tables S1-S2). But, log2FC magnitude became drastically a less accurate indicator for the direct gene targets for the later time points. This trend is expected because the downstream effects of a gene perturbation will progressively mask the true gene target identity over time. In comparison, DeltaNeTS+ prediction accuracy degraded much more mildly over the sampling times, demonstrating its higher robustness with respect to the choice of time samples (see Supplementary Tables S1-S2).

**Figure 1.**
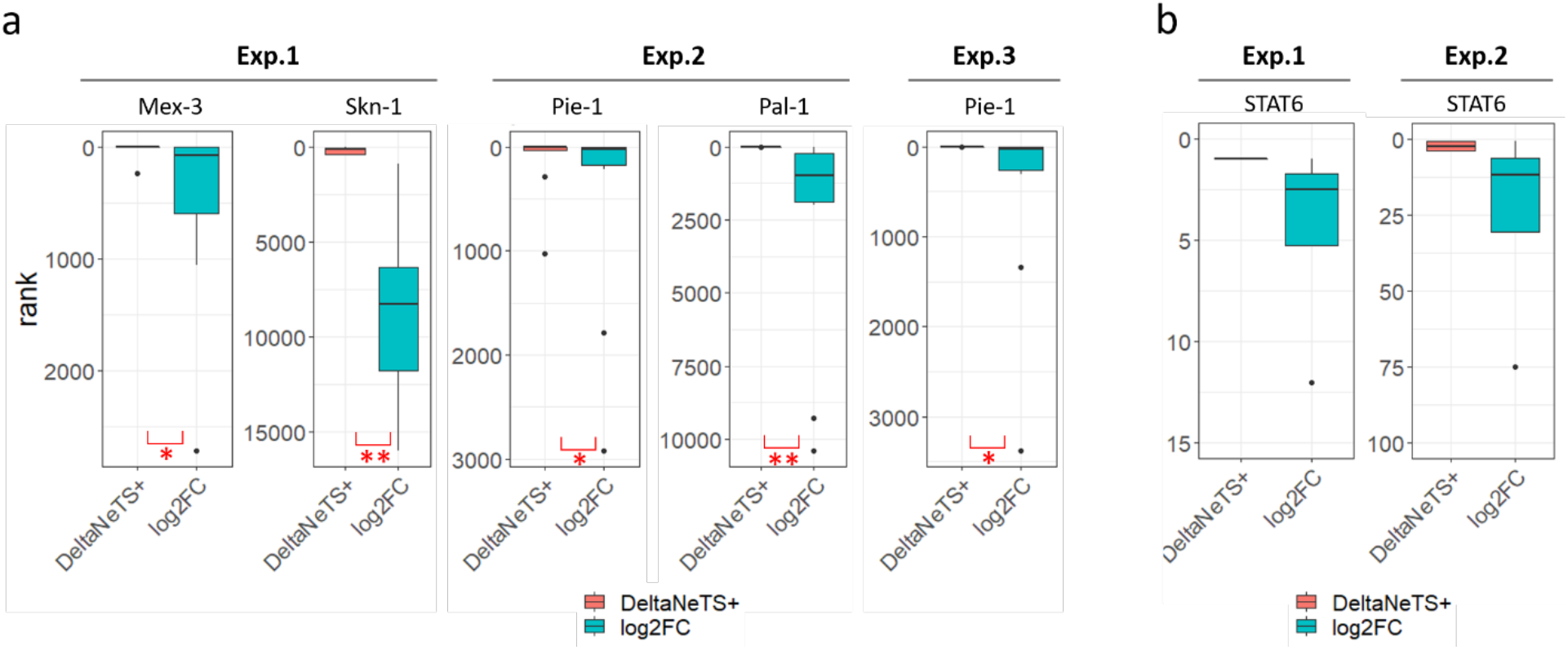
Ranks predictions for knock-out targets. The distribution of box plot indicates the ranks from all the time points from the same experiment. The ranks of gene targets by DeltaNeTS+ are colored in orange and those by log2FC analysis are colored in teal. *: p-value < 0.05 and **: p-value <0.01 by Wilcox signed rank test.

### Predicting the mechanism of interferon-α and interferon-γ actions

Next, we tested the ability of DeltaNeTS+ in differentiating the specific activity of two related compounds from the same family of proteins: interferon-α (IFN-α) and interferon-γ (IFN-γ). This task is challenging as these two signalling proteins trigger a common response of the immune system but through different signalling pathways. IFN-α and IFN-γ are cytokines that induce innate immune response against viral infection through separate signalling pathways, called type I and type II signalling, respectively. The IFN-α signal goes through type I IFN receptor and is followed by the formation of the complex IRF9, STAT1 and STAT2 that activates the transcription of IFN-α/β stimulated genes (ISG). Meanwhile, the IFN-γ signal is received by type II IFN receptor which leads to the transcription of genes containing gamma interferon activation sites (GAS) [40, 41]. The gene expression data came from a previous study using human lung cancer cells Calu-3 [31]. We employed the human epithelial lung cancer cell GRN structure as prior information for DeltaNeTS+ (see Methods). To examine cellular pathways that are perturbed by each of the two interferon treatments, we performed GO (Biological Processes) and Reactome pathway enrichment analysis for the 50 highest ranked genes from each method [35].

Table 2 summarizes the enriched GO terms and Reactome pathways of the top gene target predictions from DeltaNeTS+ and log2FC analysis (adjusted *p*-value < 0.01; detailed results in Supplementary Data 1). The top 50 gene targets predicted by TSNI did not show any statistically significant enrichment for neither GO nor Reactome pathways. The enriched GO and Reactome terms of the DeltaNeTS+ perturbation analysis correctly point to the distinct signalling pathways through which the two interferons act. Besides interferon-specific signalling, pathways related to major histocompatibility complex (MHC) class II antigen, which is induced by an IFN-γ stimulated gene CIITA [42], are also enriched in the DeltaNeTS+ analysis for IFN-γ data, but not for IFN-α data. On the other hand, the enrichment analysis of top log2FC genes shows more diverse over-represented pathways, which is expected as the log2FC expressions reflect both direct and indirect effects of the two interferons. While the distinct signalling pathways related to IFN-α and IFN-γ are among the top enriched GO and Reactome pathways for log2FC, the signalling pathways specific to each interferon are cross-listed in the enrichment results for the two interferons. In comparison to DeltaNeTS+, log2FC is less capable in differentiating the specific signalling pathways through which IFN-α and IFN-γ act.

**Table 2.**
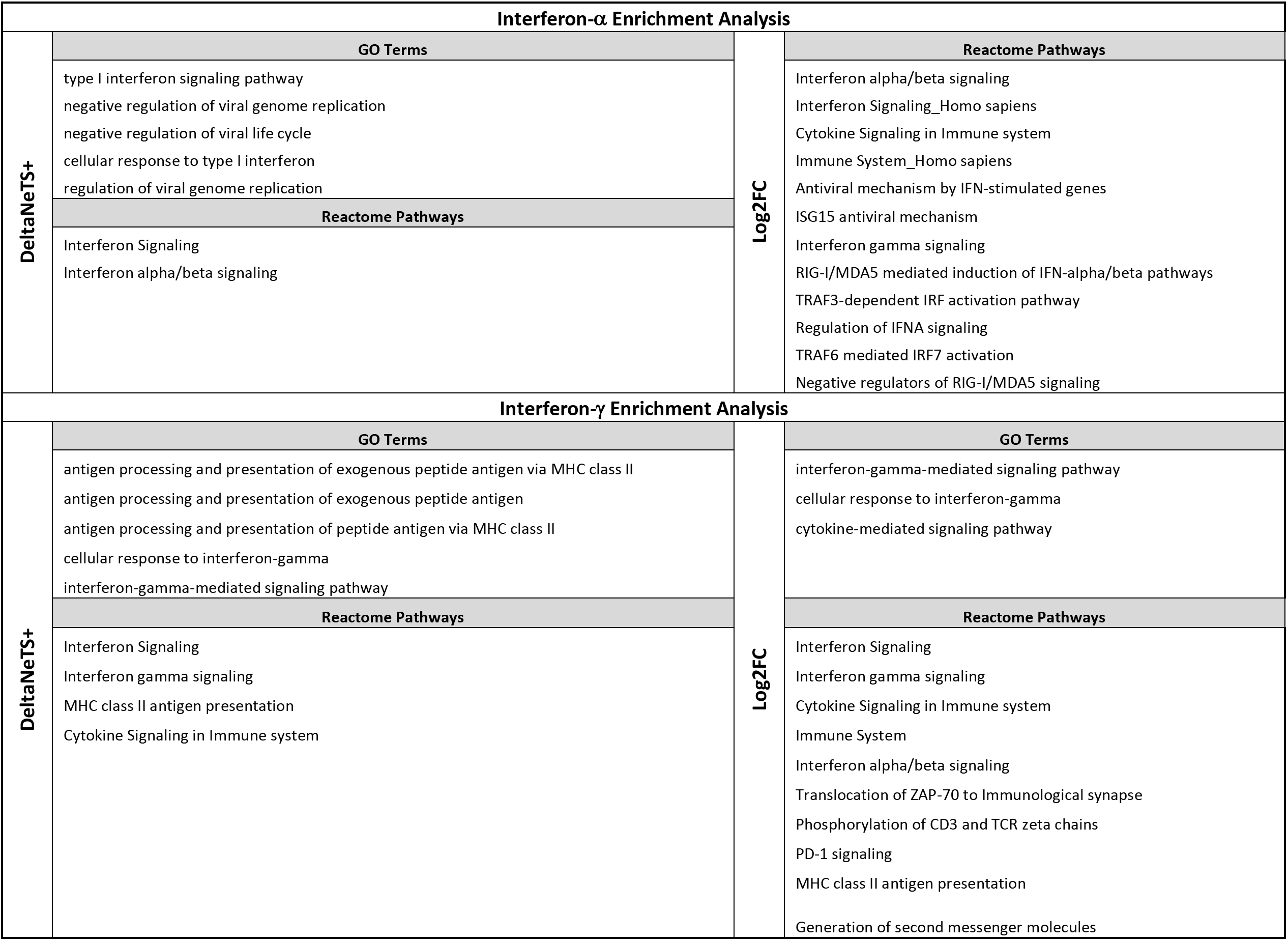
Reactome pathway enrichment analysis for Interferon-α and -γ treatments

### Network perturbation analysis during influenza viral infections

Finally, we applied DeltaNeTS+ to analyze gene expression data from H1N1 and H5N1 influenza virus infected Calu-3 cells with the goal of elucidating the similarities and differences between the two important influenza viral strains. H1N1 strain is the cause of the 2009 swine flu outbreak and is known for its high transmissibility among humans. On the other hand, H5N1 strain is an avian influenza subtype that is known for its severe virulence with a high mortality rate of 60%. This high pathogenicity results in growing attention to understand the causal molecular mechanism [43]. Following DeltaNeTS+ analysis, we performed a GSEA to find over-represented Reactome pathways [37] and WGCNA [38] analysis to identify gene modules with similar dynamic perturbations, using the gene perturbation scores produced by DeltaNeTS+.

Figure 2 depicts the Reactome pathways enriched (adjusted p-value < 0.01, see Methods) across the three phases of the infection: early (phase 1: 0-7hr), middle (phase 2: 7-18hrs) and late (phase 3: >18hrs) (full results in Supplementary Data 2). H1N1 and H5N1 trigger many of the same cellular pathways but often in different phases. Expectedly, immune response, such as cytokine signaling and innate immune system, is triggered by both viral infections with roughly the same trend over time. Other pathways however are modulated with different timings. Among the pathways significantly enriched only in H5N1 infection are those associated with G-protein coupled receptor (GPCR) signaling and actin cytoskeleton (muscle contraction), both of which are known to be hijacked for viral entry processes (Figure 2) [18, 44, 45]. On the other hand, programmed cell death and DNA damage (chromatin organization and DNA repair) are significantly enriched only in H1N1 data from the early through mid-phase of the infection. A previous study reported that death signaling molecules are downregulated in Calu-3 cells infected by H5N1 [46], a finding that is consistent with the absence of enrichment for programmed cell death in the result of DeltaNeTS+ analysis for H5N1. Other pathways enriched in the late phase of H1N1 infection are NOTCH, WNT, and RHO GTPase signaling, pathways that are relevant to cellular proliferation [47, 48]. These pathways are related to Histone cluster 1 families, and based on DeltaNeTS+ perturbation scores, are predicted to have been negatively perturbed (see Supplementary Data 3). The negative perturbations signify a repression of cellular proliferation in H1N1 infection.

**Figure 2.**
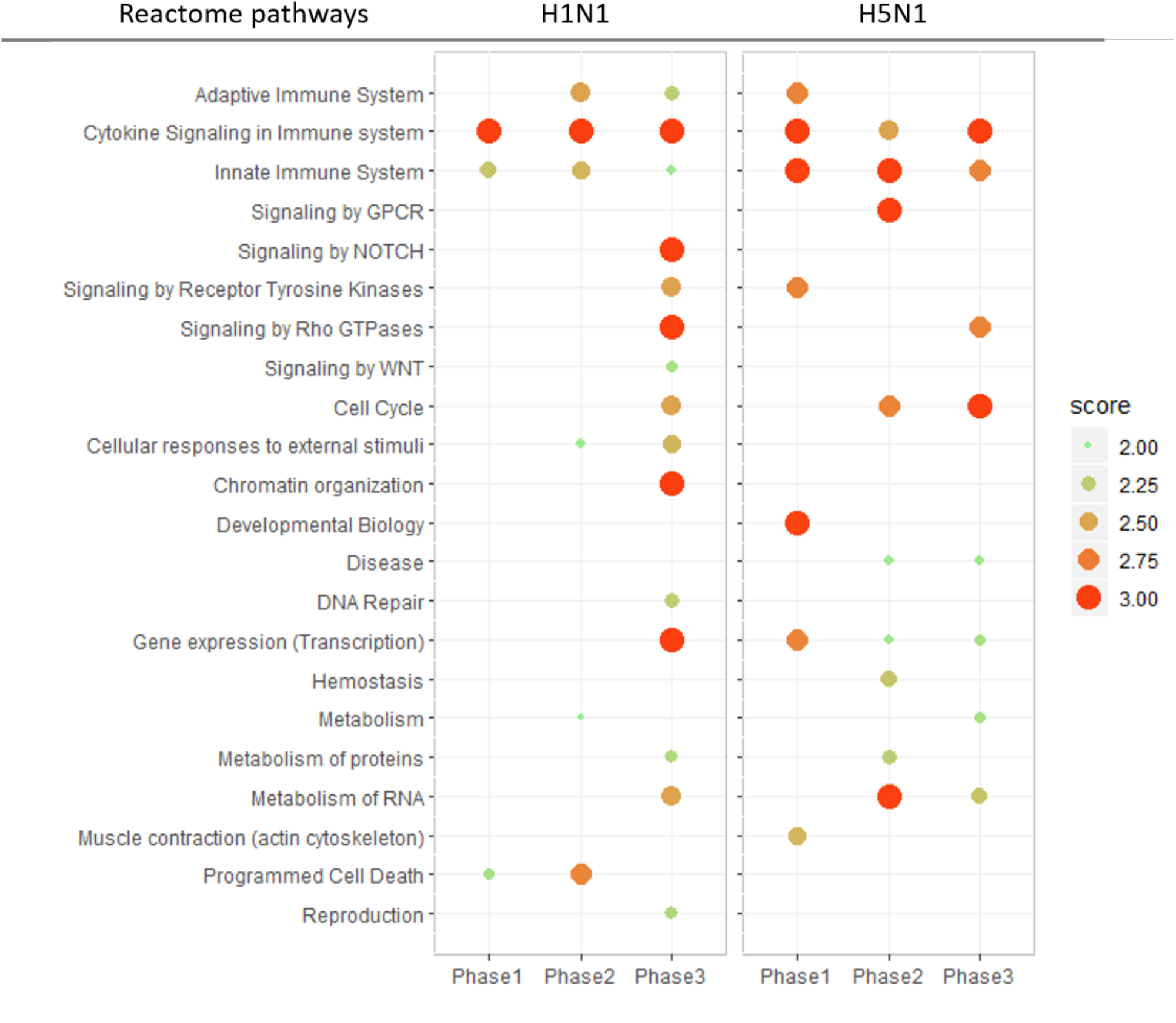
Summary of key Reactome pathways from gene set enrichment analysis of DeltaNeTS+ predictions. The size and color of dots indicate the score (negative logarithm-10 of *p*-values) for the enriched Reactome terms. The influenza infection period is divided into three time phases: Phase1 = 0-7 hours, phase 2 = 7-18 hours, and phase 3 ≥ 18 hours post-infection.

In addition to enrichment analysis, we applied WGCNA to the DeltaNeTS+ gene perturbation predictions to group the genes based on the pattern of perturbations over time. In this case, WGCNA finds clusters of genes whose perturbation scores given by the **P** matrix are highly similar based on a topological overlap derived from the gene-gene correlations. In the WGCNA results, the time-course perturbation values in each group can be represented by the first principal component in that group, called eigengene. Figure 3 shows the eigengene perturbation profiles of seven groups identified by WGCNA, and Table 3 provides the summary of GO and Reactome pathways enriched in each group (see details in Supplementary Data 4-5). For the smallest group M7, no pathway was found to be significantly enriched.

**Table 3.** WGCNA modules of DeltaNeTS+ perturbation scores

| Module | size<sup>a</sup> | Summary of enriched pathways<sup>c</sup> | DB<sup>d</sup> |
| --- | --- | --- | --- |
| M1 | 2863 | DNA replication, cell cycle | GO & RP |
| M2 | 2803 | Influenza Viral RNA Transcription and Replication, viral mRNA Translation | GO & RP |
| M3 | 2645 | Defense response to virus, immune response, interferon signaling | GO & RP |
| M4 | 1427 | Viral entry (GPCR binding, amine transport) | GO & RP |
| M5 | 888 | Crosslinking of collagen fibrils | RP |
| M6 | 762 | Apoptosis (TNFR2 non-canonical NF-κB pathway) | RP |
| M7 | 397 | N/A | - |

<sup>a</sup>The number of genes clustered in each module. <sup>b</sup> Summary of enriched pathways ( $q$ -value < 0.10) in each module. <sup>c</sup>Pathway databases used for enrichment analysis. RP: Reactome pathway database. GO: gene ontology.

**Figure 3.**
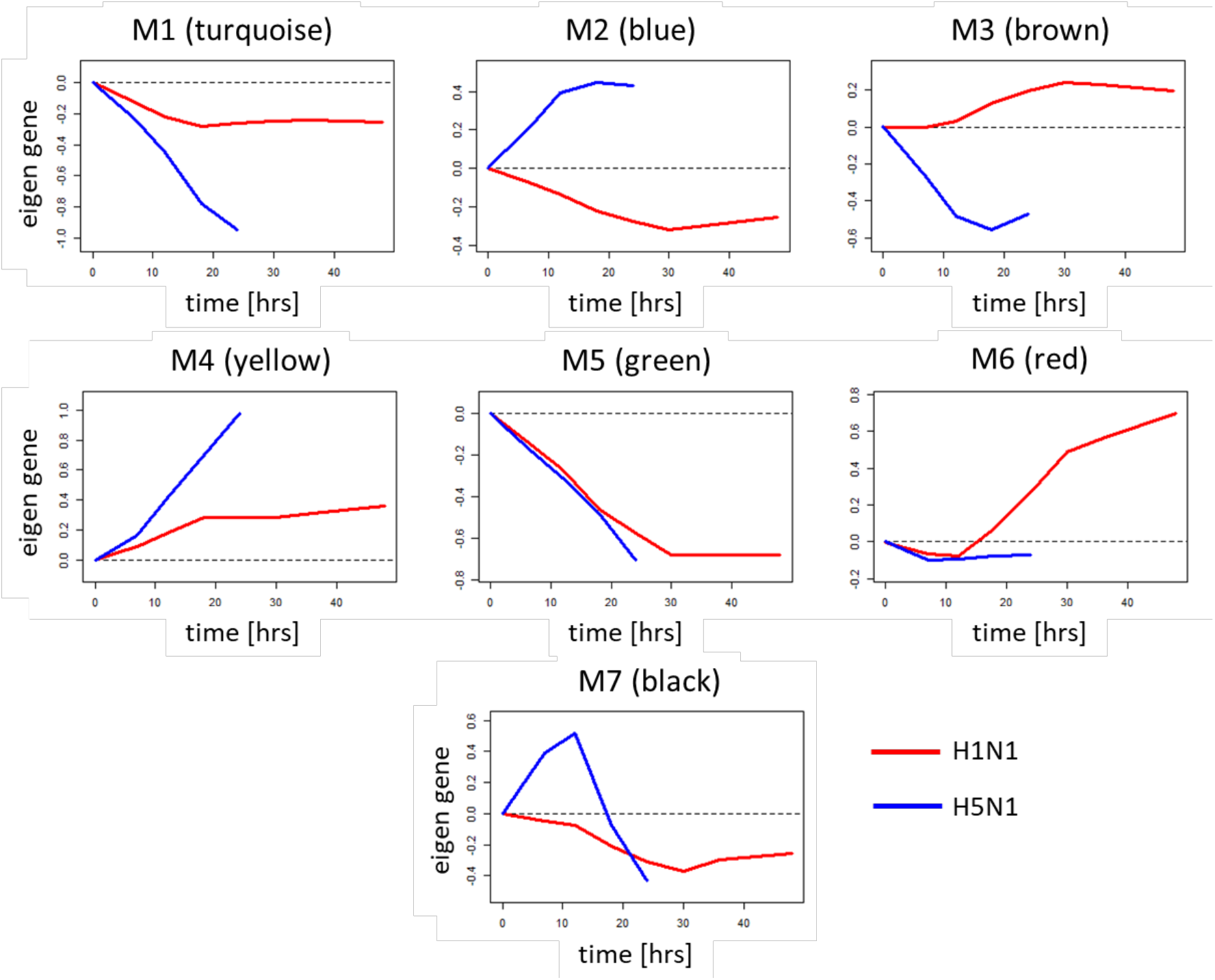
Eigengene profiles from WGCNA applied to DeltaNeTS+ perturbation scores of H1N1 and H5N1 influenza A infections (red: H1N1, blue: H5N1).

Groups M1, M4, and M5 are related to cell cycle arrest, viral entry, and cellular matrix barrier respectively. For these groups, the signs of the perturbations are the same for H1N1 and H5N1, but the magnitudes of the perturbations are larger for H5N1. For both strains, the infection acts to repress cell cycle arrest and cellular matrix barrier (negative perturbations) and to activate viral entry pathways (positive perturbations). For groups M2 and M3, the signs of the perturbations for the two influenza A strains are the opposite of each other. In H5N1 infection, viral RNA transcription, translation and replication (group M2) are induced, whereas viral defense and immune mechanisms (group M3) are strongly inhibited. In contrast, in H1N1 infection, viral RNA replication processes are repressed, but genes for antiviral mechanisms are moderately induced. Taken together, GSEA and WGCNA of DeltaNeTS+ for H1N1 and H5N1 infection in Calu3 indicates that in comparison to H1N1, the more virulent H5N1 infection shows increased activity of viral entry process and increased repression of cell cycle arrest, as well as inhibition of the immune system and programmed cell death.

## Discussion

DeltaNeTS+ is a network perturbation analysis tool for the analysis of gene expression data, which generates predictions on the direct gene expression perturbation in a given treatment or sample. DeltaNeTS+ is able to take advantage of time-series datasets of gene expression and information on the GRN structure, and thus provide accurate predictions from a small number of samples. The direct gene perturbation of a gene is computed based on the difference between the measured differential expression of the gene and the differential expression predicted by the GRN model based on the expression of its transcription factors. DeltaNeTS+ is most similar to TSNI [21], as both methods rely on a linear regression formulation and time-series data. While TSNI uses principal component analysis (PCA) to reduce the dimension of the unknown GRN matrix **A** and perturbation matrix **P**, DeltaNeTS+ employs a regularization strategy (ridge regression or LASSO, depending on the availability of the GRN structure). TSNI is unable to accommodate information on GRN structure.

Adding GRN structural information to DeltaNeTS+ is highly beneficial and improves the gene target predictions, especially when the dataset contains only a few samples. For example, DeltaNeTS+ employing LASSO applied to *C. elegans*, T-cell, and Calu-3 datasets above (number of samples < 20) gave extremely sparse solution (**P** is **0**; data not shown). Here, by using the available GRN structure, DeltaNeTS+ is implemented using ridge regression in which the dimensionality of the unknown GRN matrix **A** is reduced to known TF-gene regulations. As shown in *C. elegans* and human T-cell case study, DeltaNeTS+ was able to use the GRN structure to predict genetic perturbations with high accuracy. The ability to utilize GRN structure information is important high-quality cell type specific gene regulatory networks become more available in recent times, thanks to the success of large-scale projects such as ENCODE [17] and FANTOM5 [49]. Besides, advanced network inference tools such as ARACNE [50] and MINDy allow us to reverse-engineer a context-specific network from the gene expression data for a cell of our interest.

The application of DeltaNeTS+ to the analysis of gene perturbation targets associated with H1N1 and H5N1 infection led to insights on the differences in the mechanism of actions between the two influenza A strains. In general, the more pathogenic strain H5N1 induces stronger and swifter perturbations than the less pathogenic but more infectious H1N1 (Figures 2 and 3). Notably, H5N1 infection shows a strong induction of viral entry process and viral replication and a repression of cell cycle arrest, all of which are strategies that would ensure a successful proliferation of the virus. The pathogenic H5N1 infection also demonstrates inhibition of viral defense mechanism. Furthermore, the less severity of H1N1 infection is also associated with a successful activation of cell death and repression of cell proliferation, pathways that are crucial in curtailing viral progression.

## Conclusions

DeltaNeTS+ is an effective network analysis method for inferring gene transcriptional perturbations from gene expression dataset. The ability of DeltaNeTS+ to integrate both steady state and time-course gene expression data and available information of gene regulatory network structure enables accurate prediction of gene targets from limited number of samples (~10s). The application of DeltaNeTS+ to influenza A infection datasets in human Calu-3 give insights into the key functional perturbations that differ between highly virulent H5N1 avian flu and highly transmissible H1N1 swine flu. Insights on the molecular mechanisms of drugs and diseases from gene target predictions provide important information for drug discovery and treatment formulations.

## Ethics approval and consent to participate

Not applicable.

## Consent for publication

Not applicable.

## Availability of data and materials

DeltaNeTS+ software package is available at http://www.github.com/cabsel/deltanetsplus.

## Author’s Contributions

HN and RG designed the study. HN wrote the code. HN, JH and PC performed data analysis. HN, JES and RG interpreted the results. HN, PC, JES and RG wrote the manuscript.

## Acknowledgements

Not applicable

