## Supplementary Tables for "DeltaNeTS+: Elucidating the mechanism of drugs and diseases using gene expression and transcriptional regulatory networks"

**Supplementary Table S1.** Gene target ranking by DeltaNeTS+ and log2FC analysis for each time point in *C. elegans* datasets

| EXP1: skn-1 shRNA on mex-3 mutated cells | | | | |
| --- | --- | --- | --- | --- |
| Time [min] | mex-3 repression | | skn-1 silencing | |
|  | DeltaNeTS+ | log2FC | DeltaNeTS+ | log2FC |
| 0 | 3 | 1 | 39 | 911 |
| 23 | 4 | 2 | 46 | 6242 |
| 41 | 2 | 2 | 61 | 3889 |
| 53 | 11 | 14 | 58 | 16010 |
| 66 | 5 | 15 | 93 | 12632 |
| 83 | 7 | 137 | 210 | 13133 |
| 101 | 5 | 403 | 419 | 9266 |
| 122 | 7 | 660 | 381 | 8405 |
| 143 | 4 | 1059 | 487 | 8181 |
| 186 | 238 | 2720 | 487 | 6575 |
| EXP2: pal-1 shRNA on pie-1 mutated cells | | | | |
| Time [min] | pie-1 repression | | pal-1 silencing | |
|  | DeltaNeTS+ | log2FC | DeltaNeTS+ | log2FC |
| 0 | 2 | 1 | 1 | 2 |
| 23 | 2 | 1 | 1 | 4 |
| 41 | 4 | 2 | 1 | 1555 |
| 53 | 4 | 6 | 1 | 391 |
| 66 | 5 | 20 | 1 | 2009 |
| 83 | 8 | 32 | 1 | 195 |
| 101 | 8 | 55 | 1 | 10411 |
| 122 | 41 | 215 | 1 | 293 |
| 143 | 291 | 1787 | 2 | 9270 |
| 186 | 1029 | 2927 | 1 | 1565 |
| EXP3: pie-1 mutation | | | | |
| Time [min] | pie-1 repression | | | |
|  | DeltaNeTS+ | | log2FC | |
| 0 | 1 | | 1 | |
| 23 | 1 | | 1 | |
| 41 | 1 | | 1 | |
| 53 | 1 | | 1 | |
| 66 | 1 | | 25 | |
| 83 | 1 | | 29 | |
| 101 | 2 | | 122 | |
| 122 | 1 | | 311 | |
| 143 | 1 | | 1335 | |
| 186 | 7 | | 3385 | |

**Supplementary Table S2.** Gene target ranking by DeltaNeTS+ and log2FC analysis for each time point in STAT6 siRNA experiments of human T-cell data

|  | EXP1: STAT6 siRNA | | EXP2: STAT6 siRNA + with IL-4 | |
| --- | --- | --- | --- | --- |
| Time [h] | STAT6 silencing | | STAT6 silencing | |
|  | DeltaNeTS+ | log2FC | DeltaNeTS+ | log2FC |
| 0 | 1 | 2 | - | - |
| 12 | 1 | 1 | 1 | 2 |
| 24 | 1 | 3 | 1 | 12 |
| 48 | 1 | 2 | 2 | 92 |
| 72 | 1 | 13 | 3 | 152 |

**Supplementary Table S3.** The performances (AUROC and AUPR) of predicting gene targets in yeast data sets by DeltaNeTS with and without GRN information, DeltaNet, TSNI, and DE. The GRN information for yeast was obtained from YeasTract (Teixeira et al., 2018).

| Methods | Time series data* | | Steady-state data^§^ | |
| --- | --- | --- | --- | --- |
|  | AUROC | AUPR | AUROC | AUPR |
| DeltaNeTS + GRN info. | 0.99871 | 0.56509 | 0.95424 | 0.66388 |
| DeltaNeTS | 0.99789 | 0.79668 | 0.92754 | 0.67837 |
| DeltaNet | 0.90993 | 0.70777 | 0.76928 | 0.29214 |
| TSNI | 0.99834 | 0.53724 | - | - |
| DE | 0.99854 | 0.50334 | 0.9455 | 0.58283 |

*A yeast data set consisting of time-series gene expressions under genetic perturbations, ^§^A yeast data set consisting of steady-state gene expression profiles under genetic perturbations (see Supplementary Materials).
